# Reciprocal roles of two trehalose transporters in aestivating cabbage stem flea beetles (*Psylliodes chrysocephala*)

**DOI:** 10.1101/2025.02.22.639621

**Authors:** Gözde Güney, Doga Cedden, Stefan Scholten, Michael Rostás

## Abstract

The cabbage stem flea beetle (*Psylliodes chrysocephala*, CSFB) is a significant pest of winter oilseed rape crops in northern Europe. CSFB adults aestivate during the summer to protect themselves from heat and desiccation stress. Trehalose, the primary hemolymph sugar, has been linked to energy homeostasis and stress resilience, but its regulation and function during aestivation remain poorly understood. Here, we investigated the roles of two trehalose transporters, Tret-1 and Tret-2, in modulating trehalose dynamics across different adult stages in CSFB. Through spatiotemporal transcript profiling, we found that Tret-1 was predominantly expressed in the fat body, where it facilitates trehalose export to the hemolymph, whereas Tret-2 expression was higher in the Malpighian tubules, mediating trehalose uptake from the hemolymph. RNA interference experiments revealed that Tret-1 is involved in transporting trehalose from the fat body into the hemolymph, while Tret-2 works reciprocally to transport trehalose from the hemolymph into the Malpighian tubules. The disruption of trehalose transportation resulted in excess glucose, glycogen, and triglyceride levels, mainly in pre-aestivation beetles. Furthermore, the knockdown of either trehalose transporter caused a compensatory increase in feeding activity in pre-aestivation beetles, while the knockdown of Tret-2 compromised resilience to heat stress. Our findings uncover the reciprocal functions of Tret-1 and Tret-2 in regulating trehalose distribution and maintaining metabolic stability during aestivation, offering insights into the physiological strategies underpinning insect survival during aestivation.

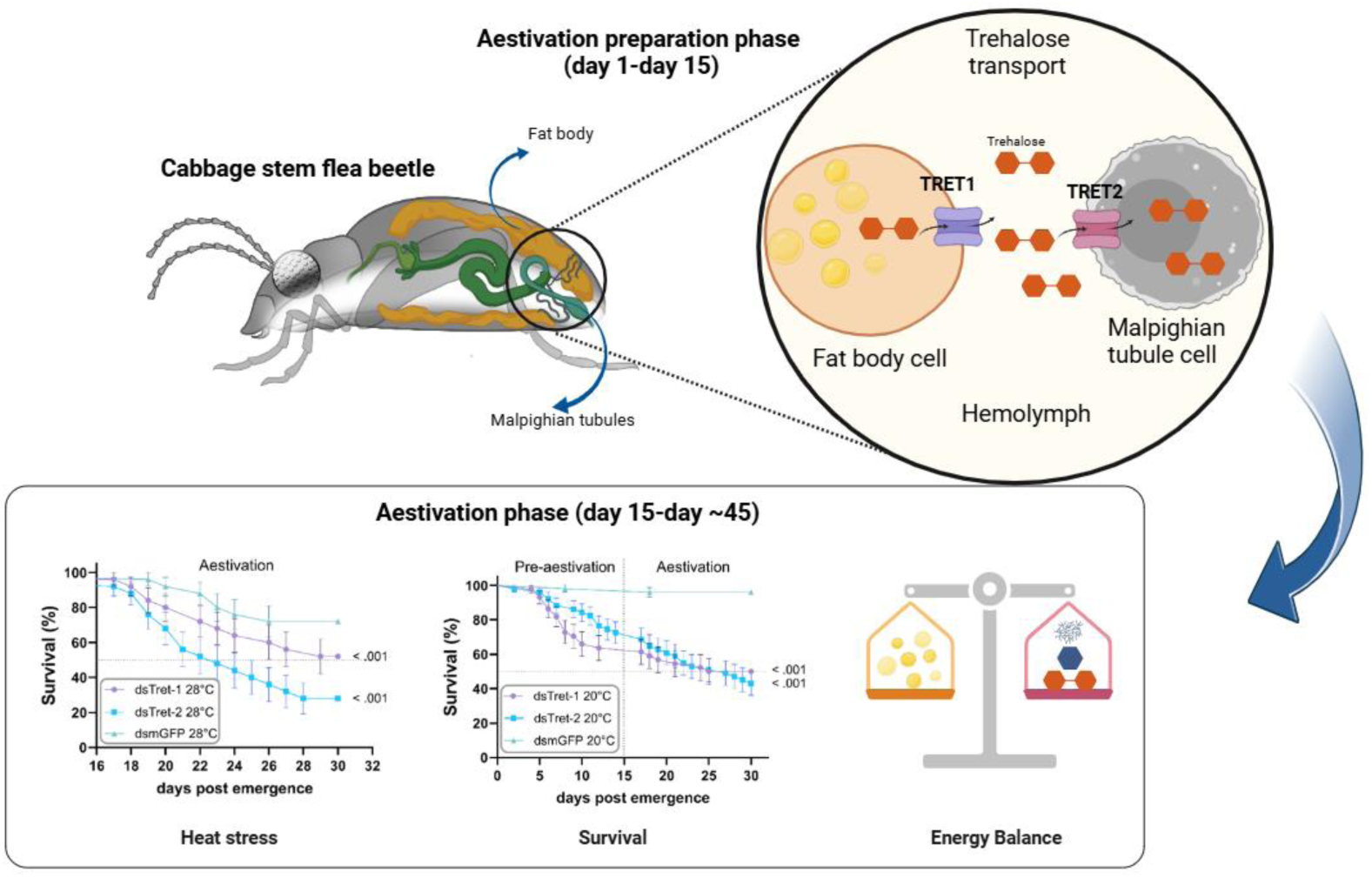

- Two Trehalose transporters were investigated in aestivating *P. chrysocephala*
- Tret-1 mainly transports trehalose out of fat body
- Tret-2 mainly transports trehalose into Malpighian tubules
- Dynamic trehalose transportation regulates other metabolites, including Glucose
- Tret-2, but not Tret-1, might be necessary for heat stress resilience

## 1 Introduction

Energy homeostasis is critical for the survival of animals. Hence, diverse metabolite transporters facilitate and direct the cellular import or export of metabolites such as trehalose (Griffiths et al., 2016). Trehalose is a nonreducing disaccharide and acts as the circulatory hemolymph sugar in insects (Becker et al., 1996; Matsushita and Nishimura, 2020; Thompson, 2003). The primary roles of trehalose include providing energy to tissues and storing long-term energy (Shukla et al., 2015). Moreover, various vital processes, such as cuticle formation and sex pheromone biosynthesis, depend on trehalose metabolism (Merzendorfer and Zimoch, 2003; Tellis et al., 2023). A less well-understood role of trehalose is its ability to serve as an intracellular bioprotectant. It has been suggested that trehalose may render cells more resilient to unfavorable environmental conditions, including heat, desiccation, and cold, by acting as a chemical chaperone (Elbein et al., 2003; Tapia and Koshland, 2014), albeit robust experimental evidence is lacking. Furthermore, the accumulation of trehalose rather than glycogen increases the lifespan of *Caenorhabditis elegans*, suggesting a general effect of trehalose on cellular viability (Seo et al., 2018).

The biosynthesis of trehalose from glucose precursors mainly occurs in the fat body, which is analogous to both the liver and adipose tissue in mammals (Becker et al., 1996; Murphy and Wyatt, 1965). The sequential reactions catalyzed by trehalose-6-phosphate synthase (TPS) and trehalose 6-phosphate phosphatase (TPP) enzymes convert glucose-6-phosphate (G6P) into trehalose (Candy and Kilby, 1961; Tellis et al., 2023). Trehalose, in turn, can be catabolized back into glucose for metabolic usage by the trehalase enzyme (Hehre et al., 1982; Shi et al., 2016; Shukla et al., 2015). As a circulatory sugar, trehalose transport into and out of specific tissues must be tightly regulated. The spatiotemporal trehalose dynamics are imposed by trehalose transporters embedded in the cellular membrane (Kanamori et al., 2010). In insects, there is more than one trehalose transporter, and its transport directionality is reversible depending on the trehalose gradient (Kikawada et al., 2007). A well-described trehalose transporter (Tret-1) in the anhydrobiotic insect *Polypedilum vanderplanki (*Diptera, Chironomidae) is dominantly expressed and accumulates trehalose in the fat body. Furthermore, its expression was increased upon salinity or desiccation stress, suggesting a role in the anhydrobiotic biology of the insect (Kikawada et al., 2007).

Diapause is a form of seasonal dormancy characterized by cessation of development in immature stages and reproductive activity in the imago stage, accompanied by reduced metabolic and behavioral activity in insects (Denlinger, 2022, 2002). Aestivation, or summer diapause, typically occurs in the summer and entails physiological changes associated with high heat stress resilience as opposed to cold resilience, as observed in hibernating (winter diapausing) insects (Saulich and Musolin, 2017). Diapause programs in most insects consist of a preparatory phase, where insects accumulate necessary metabolites and reserves, which are utilized or consumed during diapause (Denlinger, 2002; Koštál, 2006). Although there is convergence in the accumulated reserves, such as triglycerides as an energy fuel (Cedden et al., 2024c; Güney et al., 2024; Hahn and Denlinger, 2011), the accumulation of specific metabolites or proteins is dependent on the biology of the insect species, which in turn is shaped by the encountered seasonal challenges (Hahn and Denlinger, 2007). As diapause has evolved to cope with metabolic and environmental challenges, it is an excellent opportunity to comprehensively characterize the potential role of trehalose spatio-temporal dynamics in insect physiology.

The cabbage stem flea beetle (CSFB) is a significant pest of winter oilseed rape crops, especially in northern Europe. Larvae tunnel and feed inside petioles and stems, whereas adults feed on leaves and cotyledons (Ortega-Ramos et al., 2022). Managing CSFB has become challenging due to the ban on neonicotinoid seed treatments and the emergence of pyrethroid-resistant populations, the leading insecticide group used against CSFB in Europe (Højland et al., 2015; Willis et al., 2020; Zimmer et al., 2014). Hence, novel strategies, including RNA interference, are currently being developed against this pest, which requires a comprehensive understanding of pest biology at the molecular and physiological levels (Cedden et al., 2024b, 2024a). After emerging in early summer, adult *P. chrysocephala* undergo a brief period of intensive feeding before entering aestivation until August/September, usually in sheltered areas such as woodlands and hedgerows (Bonnemaison & Jourdheuil, 1954; Såringer, 1984). During aestivation, beetles do not feed or reach reproductive maturity until aestivation is terminated within 1-2 months (Ankersmit, 1964; Rüde et al., 2025; Vig, 2003). Afterward, beetles invade newly sown oilseed rape fields to feed on cotyledons or young leaves and mate (Bonnemaison & Jourdheuil, 1954; Såringer, 1984). We recently showed that aestivation in CSFB is associated with a substantial decrease in metabolism enabled by the downregulation of mitochondrial genes, increased heat tolerance, and shifts in body composition supported at the transcriptional level (Güney et al., 2024). Interestingly, trehalose transporters were among the highly differentially regulated genes between the aestivating and non-aestivating stages, raising the idea that spatiotemporal trehalose dynamics might be important for aestivation physiology.

In this study, we functionally characterized the roles of two trehalose transporters in the preparatory, aestivating, and post-aestivating CSFB. Two Trets were selected for the analysis as the transcript abundance of other Trets was very low in our previous RNA-seq study (Güney et al., 2024). First, the spatiotemporal transcript levels of the two transporters were measured alongside the concentrations of trehalose and other metabolites. Next, RNA interference was used to suppress either trehalose transporter to understand the effects on trehalose and other metabolites in different stages and tissues. Our findings suggest a model in which the reciprocal function of two trehalose transporters ensures appropriate spatiotemporal body composition dynamics, which is crucial for energy homeostasis and stress tolerance.

## 2 Materials and methods

### 2.1 Insects

The laboratory colony of cabbage stem flea beetles (*Psylliodes chrysocephala,* CSFB) was maintained at a temperature of 20° ± 1°C and a relative humidity of 65 ± 10%, with a 16:8 light: dark cycle. The beetles were reared on winter oilseed rape plants at growth stages 30–35 (BBCH scale) in rearing chambers. Freshly emerged adults were collected daily for bioassays or placed back into the rearing chambers to sustain the colony. To maintain genetic diversity, field-collected larvae from an experimental oilseed rape field in Göttingen, Germany (coordinates: 51.564065, 9.948447) were introduced annually into our laboratory colony.

### 2.2 Sequence analysis and phylogenetic tree

The putative amino acid sequences from the transcripts were obtained using ExPASy (https://web.expasy.org/translate/). Transmembrane domains were predicted using the TMHMM Server v.2.0 (http://www.cbs.dtu.dk/services/TMHMM/), while potential domains were identified using the Pfam 33.1 (http://pfam.xfam.org/) server. The NCBI accession numbers for these genes are Tret-1: GKIH01046836.1, and Tret-2: GKIH01022654.1. Phylogenetic analysis was performed using the MEGAX (https://www.megasoftware.net/) platform.

### 2.3 dsRNA synthesis

Gene-specific dsRNAs targeting coding sequence regions with no off-targets in CSFB were designed using Primer3 with default parameters. T7 promoter sequences (GAATTGTAATACGACTCACTATAGG) were added to the 5′ ends of the primer pairs (Tab. S1). Primer pairs were provided by Thermo Fisher Scientific (Germany). cDNA templates for PCR amplification were derived from RNA extracted from a mixture of ten CSFB adults aged 1–55 days, using the Quick Tissue Kit (Zymo Research, Irvine, CA, USA) and reverse transcription with the LunaScript® RT SuperMix Kit (NEB, Ipswich, MA, USA). For the synthesis of a control dsRNA, named dsmGFP, DNA template from IDT (Germany) was used. Two PCR reactions were conducted per dsRNA: one for the sense strand using T7+ forward and reverse primers and one for the antisense strand using forward and T7+ reverse primers. 50 μL PCR reactions were performed using Q5® Hot Start High-Fidelity 2X Master Mix (NEB) with 58°C annealing temperature for 30 cycles. Expected-length single bands were extracted via agarose gel electrophoresis using the Gel and PCR Clean-up Kit (Macherey-Nagel, Düren, Germany) for transcription reactions. The sense and antisense RNA strands were separately synthesized in 20 μL reactions using the MEGAscript™ T7 Transcription Kit (Invitrogen, Waltham, MA, USA) and purified by lithium chloride precipitation. Equimolar RNA strands were combined in nuclease-free water, with concentrations measured using a nanodrop (40 μg/OD260 conversion factor). The RNA strands were denatured by heating to 94°C for 5 min, then annealed at room temperature for 30 min. The length of the annealed dsRNAs was verified on a 1.5% agarose gel.

### 2.4 dsRNA feeding

The dsRNAs were diluted to 500 ng/µL in nuclease-free water and mixed with Triton-X (200 ppm, Sigma-Aldrich, St. Louis, MO, USA). Each Petri dish (60 × 15 mm with vents) contained one newly emerged adult CSFB (sex-mixed) and a 30 mm² leaf disk from the first true leaves of oilseed rape plants (BBCH ∼35), placed on 80 mm³ 1% agarose gel. 1 μL dsRNA solution was spread evenly on the leaf disk and dried for 10 min before introducing or reintroducing the beetles on days 0 (day of adult emergence), 5, and 10. Petri dishes were kept in the rearing chambers. After 24 h, when the dsRNA-treated leaf disk was consumed, the agarose gel disks were replaced with untreated 400 mm² leaf disks on agarose gel disks.

### 2.5 Body composition

Whole bodies of individual beetles were placed in 200 µL TBS (5 mM Tris [pH 6.6], 137 mM NaCl, 2.7 mM KCl), while dissected tissues (fat body, gut, muscle or Malpighian tubules) were placed in 100 µL TBS. Hemolymph was collected from decapitated beetles by centrifugation at 5,000 g for 30 min at 4°C and diluted 1:20 with TBS. All samples were frozen with liquid nitrogen and stored at −80°C. The fresh weights for the whole body and tissue samples were measured before homogenization for content normalization, while the collected hemolymph volume was measured using a pipette.

The homogenized samples were heated at 70°C for 5 min, and 20 µL of each was used to measure trehalose content using the Trehalose Assay Kit (Megazyme, Bray, Ireland), according to the manufacturer’s instructions. Additionally, 10 µL from the homogenized samples was used to measure glucose content using the D-Glucose HK Assay Kit (Megazyme) according to the manufacturer’s instructions. To determine glycogen content, 20 µL of the homogenized sample was first incubated with 80 µL of 50 mM sodium acetate buffer containing 0.5 units of amyloglycosidase (TCI, Tokyo, Japan) for 4 hours at 37°C. Following incubation, 10 µL of the sample was used to measure total glucose using the D-Glucose HK Assay Kit. Glycogen content was calculated by subtracting the glucose content from samples without amyloglucosidase treatment from the total glucose measured in the treated samples.

To measure the TAG (triglyceride) content, 2 µL of NP40 was added to 20 µL of the homogenized sample and centrifugated at 10,000 g for 10 min at 4°C. Glycerol content was determined using the Free Glycerol Reagent (Sigma) kit, and the total glycerol was measured using the Triglyceride Colorimetric Assay Kit (Cayman, Ann Arbor, MI, USA). TAG content was calculated as the difference between the total glycerol value and the glycerol content. Six to ten biological replicates were used for each measurement, with two technical replicates performed using a μQuant Universal Microplate Spectrophotometer (Bio-Tek Instruments GmbH, Bad Friedrichshall, Germany).

### 2.6 Gene expression

Whole beetle bodies or tissues (fat body, muscle, gut, or Malpighian tubules) were flash-frozen in liquid nitrogen and stored at −80°C. Total RNA was extracted using the Quick-RNA Tissue & Insect MicroPrep™ Kit (Zymo Research, Irvine, CA, USA) according to the manufacturer’s instructions. The primers were designed using the NCBI primer designing tool (https://www.ncbi.nlm.nih.gov/tools/primer-blast/). RT-qPCR was conducted with the Luna One-step RT-qPCR kit (NEB) on a 384-well plate with three technical replicates per biological replicate in a CFX384 Touch Real-Time PCR (Bio-Rad, Hercules, CA, USA). Each well contained 0.8 μL of total RNA (500 ng/μL) and 0.8 μL of a 10 mM mix of forward and reverse primers (IDT) to measure expression levels of dsRNA target genes and the reference gene rps4e, optimized for RNAi experiments (Cedden et al., 2024b). The primers targeted regions that did not overlap with the dsRNA target sequences (Tab. S1). ΔΔCq values, normalized to dsmGFP-fed samples, were used for analysis. Gene expression measurements were performed in triplicate.

### 2.7 Survival assay

Three survival experiments were conducted with three groups of beetles receiving either dsTret-1, dsTret-2, or dsmGFP. The RNAi treatments were delivered upon adult emergence. In the first survival experiment, newly emerged adults were reared under normal rearing conditions (temperature: 20° ± 1°C, relative humidity: 65 ± 10%) for 15 days post-emergence, i.e., until the onset of aestivation. In the second experiment, newly emerged adults were subjected to heat stress conditions (temperature: 28° ± 1°C, relative humidity: 50%) for 15 days post-emergence. In the third experiment, newly emerged adults were initially reared under normal rearing conditions for 15 days post-emergence, then transferred to heat stress conditions on day 16 post-emergence, where they were kept for an additional 15 days. All survival experiments replaced the agarose gel disks with untreated 400 mm² leaf disks on agarose gel disks every third day, as defined above. The survival of individual beetles (n = 30) was monitored daily.

### 2.8 Leaf consumption

As described above, newly emerged CSFBs (n = 12, mixed-sex adults) treated with 500 ng dsRNA were used to assess leaf consumption over time. The area consumed from 400 mm² leaf disks on agarose gel disks was measured on days 3, 5, 7, 9, 11, 13, and 15 post-emergence, after which feeding ceased for approximately 25 days due to aestivation. Flattened leaves were placed beside a 1 cm reference line, photographed with a 25-megapixel camera, and analyzed using ImageJ software (v1.8). The ’analyze particles’ function was used to calculate the remaining leaf area, which was subtracted from the initial 400 mm².

### 2.9 Statistical analysis

Two-way ANOVA followed by Tukey’s test was used to analyze gene expression and trehalose, glucose, glycogen, and triglyceride content data. Two-way ANOVA followed by Dunnett’s test (control: dsmGFP group) was used to analyze leaf consumption data. Kaplan–Meier survival curves were plotted using the results from the survival experiments. Curves of dsRNA treatment groups were compared to the dsmGFP group using pairwise log-rank tests with Bonferroni correction for each experiment. Results were considered statistically significant at *P* < 0.05. All statistical procedures were conducted using GraphPad Prism v10.1.

## 3 Results

### 3.1 Characterization of Tret-1 and Tret-2

Sequence analysis showed that Tret-1 ORF is 1,524 bp long and encodes 508 amino acids, while the Tret-2 ORF has a size of 1,404 bp, encoding 468 amino acids. The predicted protein weights are 56.47 kDa for Tret-1 and 50.89 kDa for Tret-2. Domain analysis for both TRET-1 and TRET-2 proteins showed the presence of 12 transmembrane domains. Additionally, domain analyses indicated that both proteins belong to the major facilitator superfamily (MFS) and contain sugar transporter-like and trehalose transporter domains. However, TRET-2, unlike TRET-1, also possesses a sugar/inositol transporter domain (Fig. 1a). *P. chrysocephala* TRET-1 and TRET-2 proteins are classified within the Coleoptera branch in the phylogenetic tree (Fig. 1b). An RT-qPCR experiment was conducted to measure the transcript levels of Tret-1 and Tret-2 at pre-aestivation (5 day old adult), aestivation (30 day old adult), and post-aestivation (55 day old adult) stages in whole beetle bodies. Tret-1 transcript levels were highest at the pre-aestivation stage and declined at both the aestivation and post-aestivation stages. Although Tret-2 transcript levels also peaked during the pre-aestivation phase, its abundance did not significantly decrease at the aestivation stage. However, similar to Tret-1, Tret-2 transcript levels declined at the post-aestivation stage (Fig. 1c, d). To further investigate tissue-specific expression, we measured the transcript levels of Tret-1 and Tret-2 separately in the fat body, Malpighian tubules, gut, and muscle of pre-aestivation beetles. The results showed that Tret-1 was abundantly expressed in the fat body, whereas Tret-2 transcript levels peaked in the Malpighian tubules (Fig. 1e).

**Figure 1.**
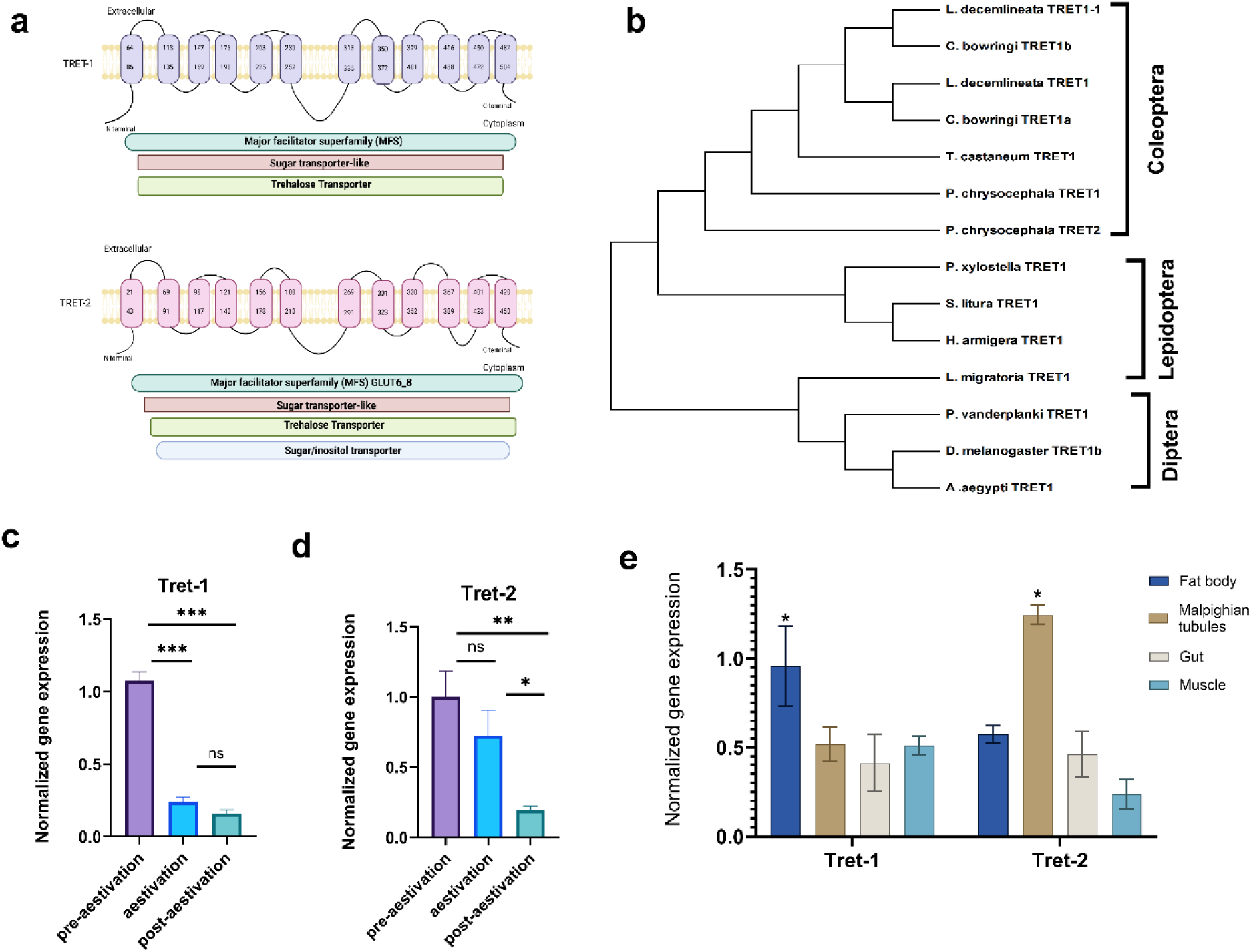
Putative protein domains, phylogenetic tree, and spatiotemporal expression dynamics of trehalose transporters. (a) The positions of the transmembrane helices and domain boundaries are labeled with the corresponding amino acid numbers. The functional domains for both proteins are the major facilitator superfamily (MFS) domain, the sugar transporter-like domain, and the trehalose transporter domain. For TRET-2, an additional sugar/inositol transporter domain was present. (b) Phylogenetic tree of *P. chrysocephala* TRET-1 and TRET-2 proteins and other 12 TRET proteins from different insects: *Leptinotarsa decemlineata* TRET1-1 (XP_023020580.1), *Colaphellus bowringi* (TRET1a, MN971589; TRET1b, MN971590), *Tribolium castaneum* TRET1 (XP_064214892.1), *Locusta migratoria* TRET1 (AAT72921.1), *Polypedilum vanderplanki* TRET1 (KAG5676724.1), *Aedes aegypti* (XP_021698361.1), *Drosophila melanogaster* TRET1b (NP_725068.1), *Plutella xylostella* TRET1 (XP_011568338.1), *Helicoverpa armigera* TRET1 (XP_021195453.3), *Spodoptera litura* TRET1 (XP_022836377.1). Expression levels of Tret-1 (c) and Tret-2 (d) at pre-aestivation, aestivation, and post-aestivation stages (n = 3). Statistical analysis was performed using one-way ANOVA followed by Tukey’s test to compare the three stages, with significant differences indicated by ***p < 0.001, **p < 0.01, *p < 0.05, n.s. = not significant. (c) Tissue-specific expression of Tret-1 and Tret-2 in the fat body, Malpighian tubule, gut, and muscle. ANOVA followed by Tukey’s test was conducted to compare the four tissues, with significant differences indicated by (*p < 0.05).

### 3.2 Spatio-temporal trehalose dynamics

Next, the trehalose content was investigated in the fat body, hemolymph, and Malpighian tubules following adult emergence. The trehalose content in the fat body was highest at the first time point after emergence and steadily declined until day 55 (Fig. 2a). In contrast, the trehalose content in hemolymph steadily increased until mid-aestivation (day 30) followed by a decline by day 55 (Fig. 2b). The pattern in Malpighian tubules was similar to that in hemolymph with higher abundance in days 18 and 30 (both within aestivation period) with a decline by day 55 (Fig. 2c).

**Figure 2.**
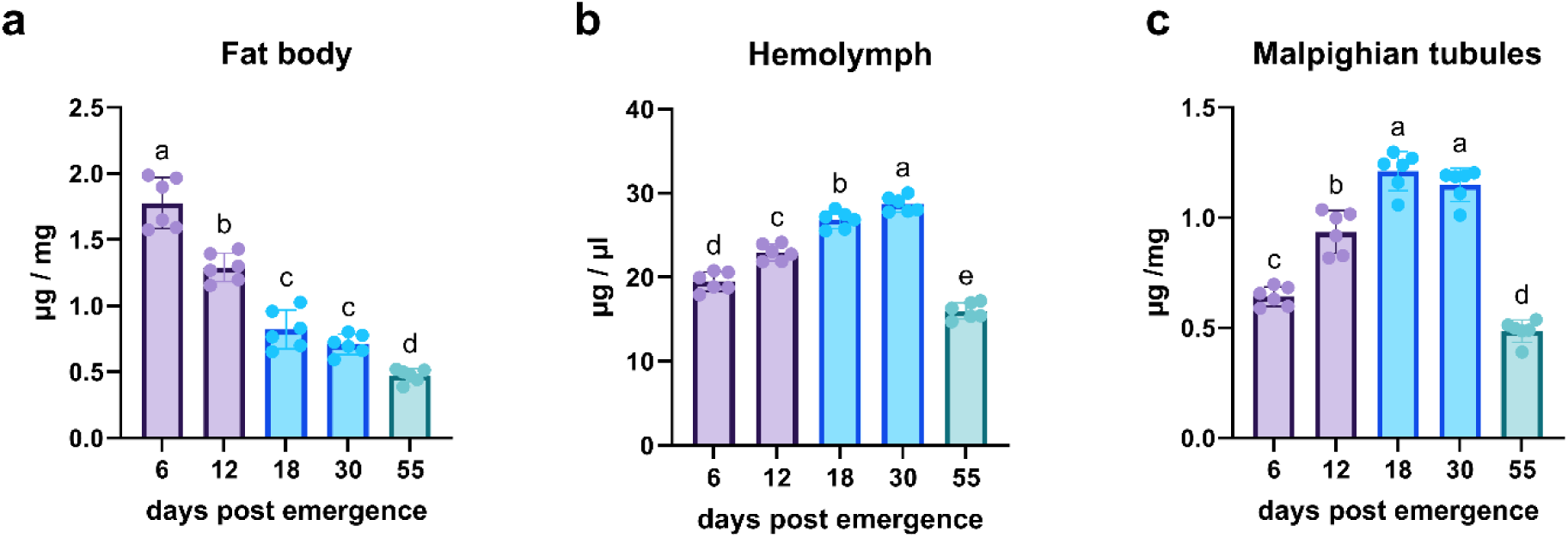
Spatio-temporal trehalose dynamics. Trehalose content in the fat body (a), hemolymph (b), and Malpighian tubules (c). Different letters above the bars indicate significant differences between the time points as determined by ANOVA followed by Tukey’s multiple comparison test (p < 0.05) (n = 6-10).

### 3.3 RNAi of Tret-1 and Tret-2

To functionally characterize the two Trets, we performed RNAi experiments by orally administering gene-specific dsRNAs, following the method established by Cedden et al. (2024b). The results demonstrated that dsRNA treatments significantly reduced the transcript abundance of the targeted Trets in whole bodies on days 5 and 15. Interestingly, silencing Tret-2 also led to a slight reduction in Tret-1 transcript levels, whereas knockdown of Tret-1 increased Tret-2 transcript abundance (Fig. 3a, b).

**Figure 3.**
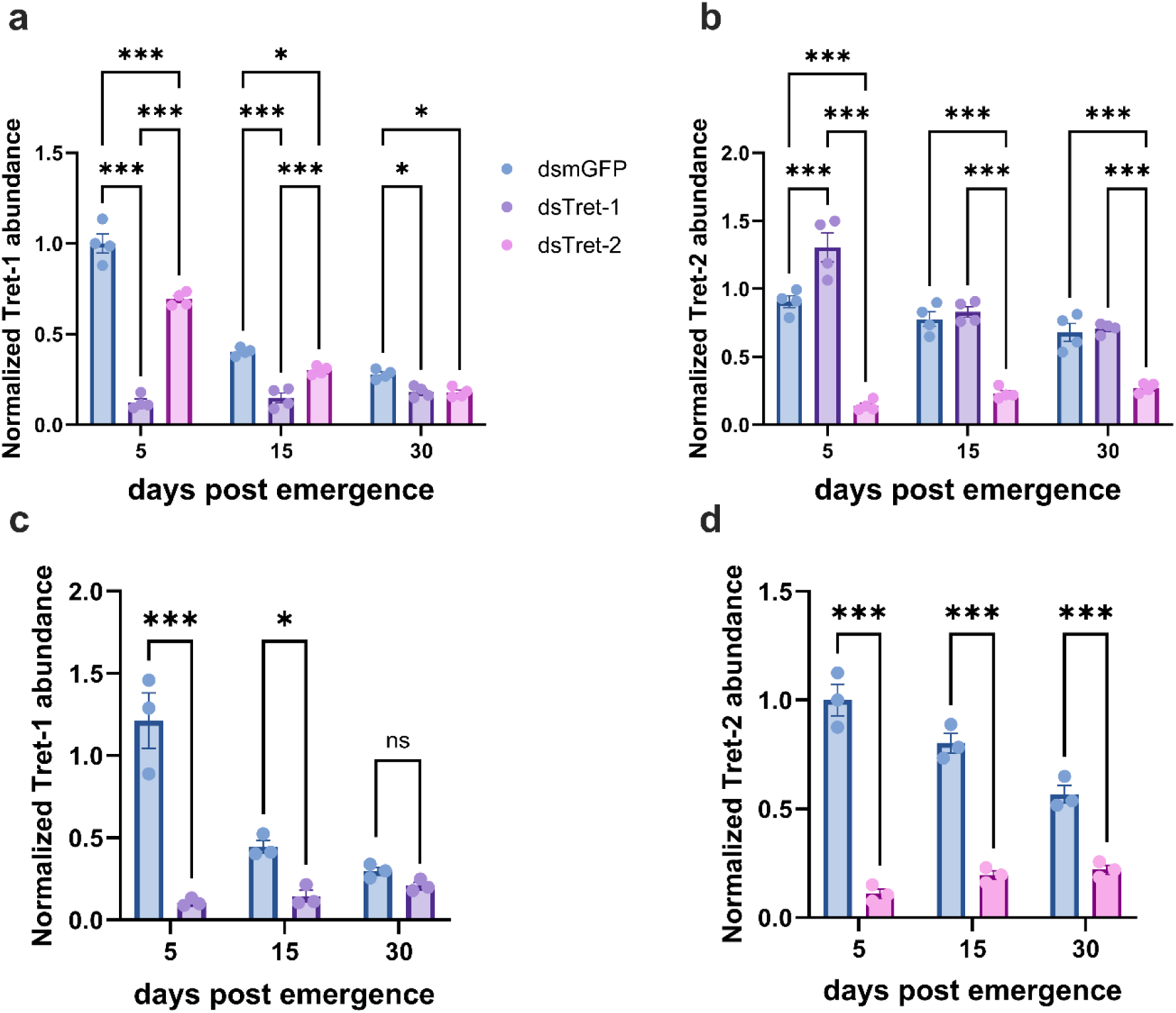
RNAi of Tret-1 and Tret-2. Gene expression levels were measured on days 5 (pre-aestivation), 15 (early-aestivation), and 30 (mid-aestivation) following dsRNA treatment of beetles. (a) Expression levels of Tret-1 in the whole body when treated with dsTret-1 dsTret-2 or dsmGFP (b) Expression of Tret-2 in the whole body when treated with dsTret-1, dsTret-2 or dsmGFP. Tret-1 (c) and Tret-2 (d) expression levels in the fat body and Malpighian tubules after treatment with dsTret-1, Tret-2, or dsmGFP as control. Statistical analysis was performed using two-way ANOVA followed by Tukey’s test, with significant differences indicated by ***p < 0.001, **p < 0.01, *p < 0.05, n.s. = not significant, respectively (n = 3-4).

We also measured the transcript abundance of Tret-1 and Tret-2 in the fat body and Malpighian tubules, i.e., the tissues with the highest abundance (Fig. 3c, d). The results were similar to whole-body measurements where RNAi of Tret-1 significantly reduced the target transcript levels on days 5 and 15 but not on day 30, and RNAi of Tret-2 significantly reduced the target transcript on days 5, 15, and 30. These measurements confirmed that RNAi treatments achieved the expected knockdown of their targets in the tissues where they are dominantly expressed.

### 3.4 Spatio-temporal dynamics of trehalose content following RNAi

Following confirmation of RNAi knockdown of both Trets, we measured spatiotemporal trehalose concentrations (Fig. 4a-c). Notably, RNAi of Tret-1 resulted in a twofold increase in the concentration of trehalose in the fat body on day 5, though this effect diminished at later time points. RNAi of Tret-2 caused a minor but significant increase in trehalose concentrations in the fat body on days 5 and 15. In contrast, the effect of RNAi on Tret-1 was reversed in the hemolymph, where trehalose levels decreased on days 5, 15, and 30. Conversely, silencing Tret-2 increased trehalose levels in the hemolymph. In the Malpighian tubules, RNAi of both Trets led to similar effects. Silencing Tret-1 decreased trehalose levels on days 15 and 30 while silencing Tret-2 decreased trehalose levels on days 5, 15, and 30. RNAi of Tret-2 also caused a decrease in transcript levels of Tret-1 on days 5, 15, and 30, albeit to a much lower degree.

**Figure 4.**
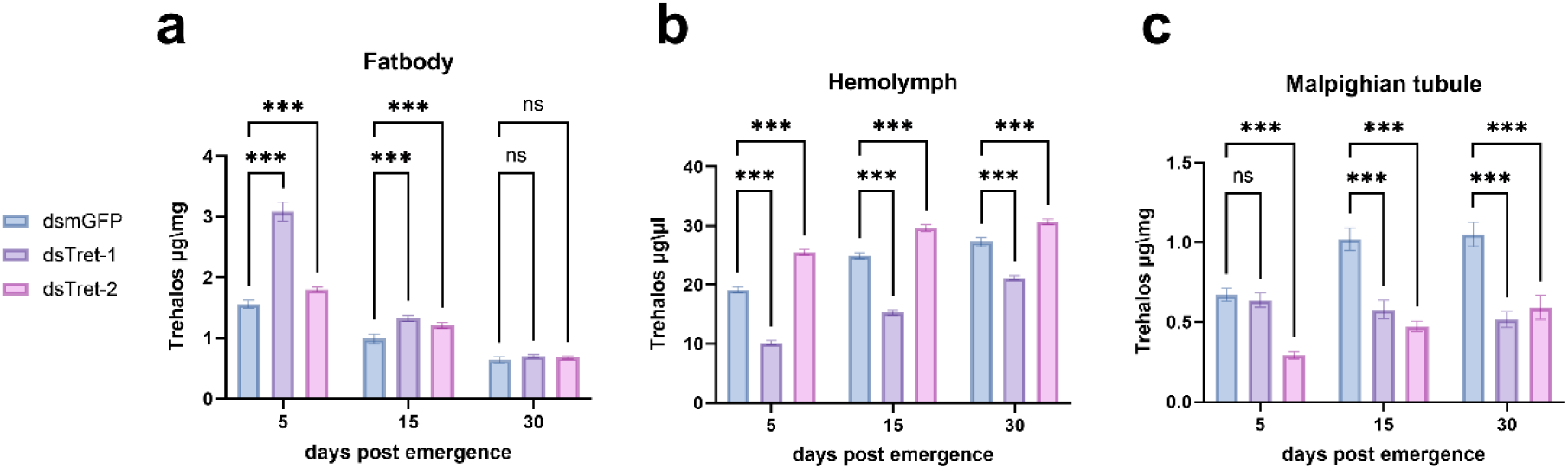
Spatio-temporal dynamics of trehalose content following dsRNA feeding. Trehalose concentrations were measured in the fat body (a), hemolymph (b), and Malpighian tubules (c) after treatment with dsTret-1, dsTret-2, and dsmGFP on days 5 (pre-aestivation), 15 (early-aestivation), and 30 (mid-aestivation). Statistical analysis was performed using two-way ANOVA followed by Tukey’s test to compare trehalose concentrations across treatments and time points. Significant differences between treatment groups are indicated by ***p < 0.001, **p < 0.01, *p < 0.05, n.s. = not significant, respectively (n = 6-10).

### 3.5 Body composition measurements

Next, we measured the trehalose, glucose, glycogen, and TAG levels in the whole body, which could be influenced by the impaired trehalose transport following RNAi of the two Trets. Trehalose levels increased in the whole body on day 5 after RNAi of either Tret-1 or Tret-2 (Fig. 5a). This difference diminished by day 15 in the Tret-1 RNAi group and by day 30 in the Tret-2 RNAi group. Glucose levels increased on day 5 following RNAi of either Tret, but this effect persisted only in the Tret-2 RNAi group on day 15 (Fig. 5b). Similarly, glycogen levels increased on days 5 and 15 after RNAi of either Tret, with a sustained increase only in the Tret-2 RNAi group on day 30 (Fig. 5c). TAG levels also increased on day 5 following RNAi of either Tret, though to a lesser extent. However, only RNAi of Tret-1 led to a significant increase in TAG levels on day 15 (Fig. 5d). These results indicate that impaired trehalose transport leads to significant changes in the concentrations of other primary metabolites.

**Figure 5.**
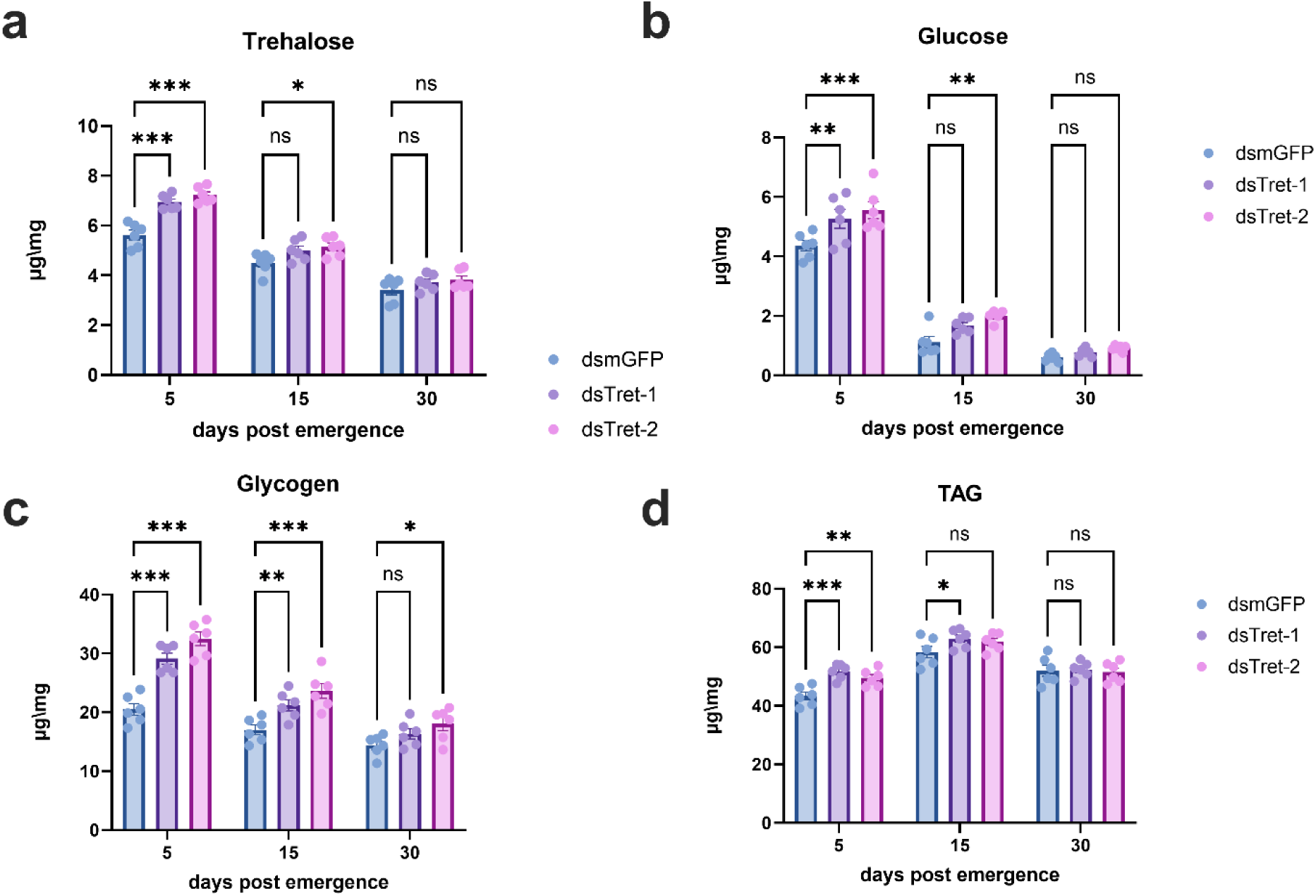
Whole body metabolite concentrations in CSFB at days 5 (pre-aestivation), 15 (early-aestivation), and 30 (mid-aestivation) following dsRNA treatments. Concentrations of trehalose (a), glucose (b), glycogen (c), and triglycerides (TAG, d) were measured after treatment with dsTret-1, dsTret-2, and dsmGFP. Statistical analysis was performed using two-way ANOVA followed by Dunnett’s multiple comparison test to compare metabolite concentrations across different treatments and time points. Significant differences are indicated by ***p < 0.001, **p < 0.01, *p < 0.05, n.s. = not significant, respectively (n = 6-10).

### 3.6 Effects of RNAi treatments on feeding behavior and survival

The observation that compromised trehalose transport caused alterations in trehalose, and other major metabolite concentrations prompted us to investigate whether silencing of Trets would affect the feeding behavior and heat stress tolerance of CSFB. Temporal feeding measurements showed that beetles fed significantly more following RNAi of Tret-2 on all days except day 15, when they naturally cease feeding activity upon entering aestivation. Similarly, RNAi of Tret-1 also increased feeding activity at most time points (Fig. 6a).

**Figure 6.**
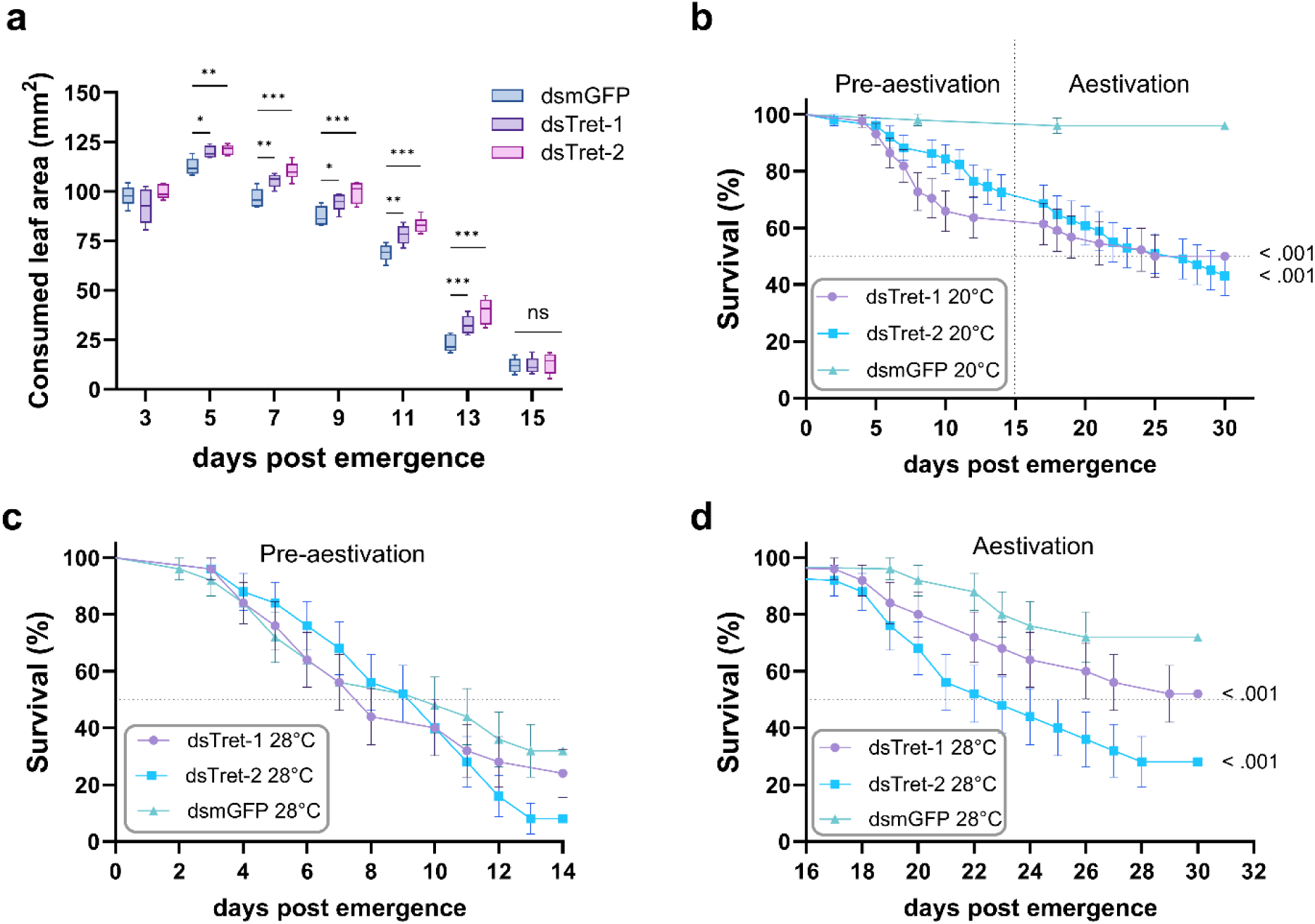
Effects of dsRNA treatments on feeding behavior and survival. (a) Consumed leaf area (mm²) measured over 15 days post-emergence in dsTret-1, dsTret-2, and dsmGFP treatment groups. Beetles were fed 500 ng dsRNA on days 0 (adult emergence), 5, and 10. Statistical analysis was performed using two-way ANOVA followed by Dunnett’s test, with significant differences indicated by ***p < 0.001, **p < 0.01, *p < 0.05, n.s. = not significant, respectively (n = 12). (b-d) Survival analyses for dsTret-1, dsTret-2, and dsmGFP groups. (b) Newly emerged CSFB adults received one of the dsRNA treatments and were reared at 20 C° for 30 days, with days 0-14 representing the pre-aestivation period and days 15-30 representing the first half of aestivation. (c) Newly emerged CSFB adults were treated with dsRNA and reared at 28 C° until day 15 (aestivation onset). (d) CSFB adults were treated with dsRNA and reared at 28 C° during the first half of aestivation. Statistical significance for survival differences was determined using the Logrank test, with significance indicated (p < 0.05) (n = 30).

To investigate the effects of RNAi-mediated silencing of Trets on survival under heat stress, we first examined baseline survival under normal rearing conditions (20 °C) immediately after adult emergence (Fig. 6b). RNAi of either Tret significantly reduced the survival of the beetle population by around 50% compared to around 95% in the dsmGFP control group under normal conditions. When beetles were reared at 28 °C (heat stress) after emergence, the survival of both gene-silenced and control groups dropped below 40% with no significant difference between groups (Fig. 6c). In a second survival experiment, beetles were initially reared under normal conditions following adult emergence and received one of the RNAi treatments. However, after day 15 (the onset of aestivation), they were maintained at 28 °C (Fig. 6d). This experiment aimed to determine whether RNAi of Trets affects the heat stress tolerance typically conferred by aestivation physiology. Interestingly, while the control group exhibited approximately 70% survival by day 30, RNAi of Tret-2 significantly reduced survival to around 30%, and RNAi of Tret-1 lowered survival to approximately 50%—a level comparable to the survival observed under normal conditions with the same RNAi treatment.

## 4 Discussion

The cabbage stem flea beetle (CSFB) is a major pest of oilseed rape plants (Döring and Ulber, 2020; Zheng et al., 2020) and has an interesting life cycle that includes obligatory aestivation in the summer (Såringer, 1984). Our previous transcriptomics study suggested that changes in trehalose transport might be important in the physiological regulation of aestivation (Güney et al., 2024). In the present study, we characterized the functional roles of two trehalose transporters in pre-aestivating and aestivating CSFBs. Our findings suggest a model in which the reciprocal function of the two trehalose transporters ensures appropriate spatiotemporal trehalose dynamics, which are crucial for energy homeostasis and stress tolerance.

The spatiotemporal measurements in this study showed that trehalose levels gradually decline in the fat body following adult emergence, while trehalose levels peak in the hemolymph and Malpighian tubules during aestivation. However, in the post-aestivation, trehalose levels decreased in the fat body, hemolymph, and Malpighian tubules. Similarly, in other insects such as *Delia antiqua*, trehalose levels diminish following the termination of diapause (Guo et al., 2015), and trehalase enzyme is upregulated in the post-diapause stage of *Antheraea pernyi* to decrease in hemolymph trehalose levels (Y.-N. Li et al., 2020). Furthermore, reproductively active *Colaphellus bowringi* females transport more trehalose into the ovaries than diapausing females (J.-X. Li et al., 2020). Although this study does not focus on the ovaries, it suggests that trehalose may be an energy source for reproduction during the post-diapause phase in CSFB. The gradual decrease in trehalose levels in the fat body of CSFB indicates that the high levels observed in pre-aestivation may not be essential. Instead, trehalose appears to be progressively utilized to sustain the beetles through the aestivation and post-aestivation stages. During pre-aestivation, the high trehalose levels might be necessary to accumulate other long-term energy stores such as TAGs, which occur primarily in the fat body. In Colorado potato beetles (CPB), adults in the pre-diapausing stage accumulate high amounts of TAGs through *de novo* lipogenesis (Cedden et al., 2024c). *De novo* lipogenesis is primarily driven by sugar metabolites (Doğan et al., 2021; Saraiva et al., 2021), which can be rapidly provided by trehalose metabolism regulated by the Trets in CSFB. The trehalose content in the hemolymph and Malpighian tubules steadily increased until mid-aestivation (day 30), then declined by day 55. This provides evidence that the trehalose abundantly accumulated in the fat body of pre-aestivation CSFB is gradually released into the hemolymph, which is likely used as both a fuel source and a bioprotectant. As aestivating CSFBs do not feed, the observed spatiotemporal changes in trehalose levels before and during aestivation are driven by trehalose transportation dynamics. Similar gradual releases of metabolites such as TAG and glycogen to maintain energy homeostasis during diapause have been observed in diverse insects, including the CPB (King et al., 2020; Lefevere et al., 1989). The gradual mobilization of these energy stores has been attributed to hormonal regulation, particularly by adipokinetic hormone (Lorenz and Gäde, 2009; Toprak, 2020; Van der Horst et al., 2001). Further research is needed to identify the endocrine or other signaling mechanisms that regulate trehalose transporters in CSFB, as demonstrated in this study, or to uncover additional yet unknown regulatory pathways.

Our results confirm the dominant role of Tret-1 in the fat body, suggesting that it functions to export trehalose to the hemolymph as its suppression increased trehalose levels in the fat body while decreasing it in the hemolymph. The same phenotype was observed in *Anopheles gambiae*, where RNAi-mediated knockdown of AgTret1 decreased hemolymph trehalose concentration by 40% (Liu et al., 2013). The results also suggest that Tret-2 is involved in importing trehalose from the hemolymph into the Malpighian tubules, as its suppression led to increased trehalose levels in the hemolymph while decreasing them in the Malpighian tubules. Although RNAi of Tret-1 also decreased trehalose levels in the Malpighian tubules, this could be conversely attributed to an indirect effect mediated by decreased trehalose levels in the hemolymph. Trehalose transporters have been described as reversible in direction (Kikawada et al., 2007), which aligns with the complex spatiotemporal trehalose regulation suggested by our findings in CSFB. Trehalose transport into the Malpighian tubules by Tret-2 may require low pH conditions to facilitate the proton-dependent process, as observed in the brown planthopper (Kikuta et al., 2012).

Suppression of Tret-1 trapped a high amount of trehalose in the fat body, which would cause the fat body cells to cease trehalose production due to intracellular trehalose saturation. Likewise, silencing Tret-1 in *C. bowringi* resulted in increased trehalose levels in the fat body of diapause-destined beetles (J.-X. Li et al., 2020), highlighting the conserved function of Tret in exporting trehalose into the hemolymph during diapause programs. The accumulation of trehalose in the fat body would, in turn, elevate cellular glucose levels, as glucose — specifically glucose 6-phosphate and UDP-glucose — is the primary precursor for trehalose synthesis in the fat body (Murphy and Wyatt, 1965). The increase in glucose levels also explains the subsequent increase in its polymeric form, glycogen. The slight increase in TAG levels following RNAi of Tret-1 could also be explained by a similar mechanism, where the accumulation of excess energy metabolites stimulates the production of TAGs, likely via *de novo* lipogenesis in the fat body (Cedden et al., 2024c). In contrast, RNAi of Tret-2 mainly increased trehalose levels in the hemolymph. Hence, this increase in energy content is likely due to increased uptake of excess trehalose by other tissues that were not investigated in the study, such as muscle, along with the slight increase in trehalose in the fat body, which is probably caused by the inhibition of trehalose uptake by the Malpighian tubules.

RNAi of Tret-2 caused downregulation of Tret-1, which is likely not a direct effect of the RNAi (since the dsRNAs do not cross-target each other due to a lack of complementarity, checked via dsRIP, https://dsrip.uni-goettingen.de/). Rather, we argue that the increased trehalose levels in the hemolymph following RNAi of Tret-2 signaled to the fat body that Tret-1 is less required to maintain trehalose homeostasis. Interestingly, the opposite effect was observed following RNAi of Tret-1, which increased Tret-2 transcript abundance on days 5, 15, and 30. This suggests a compensatory mechanism in which the suppression of Tret-1 and the consequent decrease in hemolymph trehalose levels triggers upregulation of Tret-2 to ensure sufficient trehalose transport into the Malpighian tubules. Further work should focus on the potential cross-talk mechanisms responsible for these compensatory gene regulations. For instance, cross-talk between the fat body and the brain, mediated by circulating Krebs cycle metabolites in the hemolymph, regulates diapause termination (Xu et al., 2012). Additionally, the transport of trehalose into the ovaries of *Rhodnius prolixus* is regulated by ILP, and AKH signaling cascades through the regulation of Tret expression (Leyria et al., 2021). Depending on spatiotemporal trehalose levels, similar mechanisms may regulate the expression of different trehalose transporters across tissues.

Both Trets are essential genes, with RNAi of either leading to significant mortality under normal conditions, making it difficult to solely attribute the reduced survival under heat stress to decreased resilience. Nevertheless, our findings suggest that Tret-2, but likely not Tret-1, plays a role in fine-tuning spatiotemporal trehalose levels to increase heat stress tolerance. Under heat stress, insects rapidly lose water, increasing the risk of desiccation (Benoit, 2010). We propose that desiccation is the primary driver of mortality in our experimental setup, as 28 °C is not extreme enough to cause death by itself. In *A. gambiae*, RNAi of Tret-1 (a fat body Tret) decreased resilience to desiccation and heat (Liu et al., 2013), suggesting that different Trets play distinct roles in heat and desiccation resilience across species. In the case of CSFB, Tret-2 (a Malpighian tubule Tret) appears to be critical for conferring this resilience. Similarly, the knockdown of a Tret-1-like gene in *Plutella xylostella* reduced survival under heat stress in addition to reducing fecundity (Zhou et al., 2022). Trehalose-conferred resilience to heat or desiccation stress was also observed in the yeast model *Saccharomyces cerevisiae*, where intracellular trehalose enhances resilience against desiccation rather than supporting metabolism (Tapia et al., 2015). However, trehalose has also been suggested to serve as an energy source that supports recovery in anhydrobiotic insects following complete desiccation (Ryabova et al., 2020). Hence, Tret-2 is likely contributing to desiccation resilience by maintaining appropriate intracellular trehalose levels, albeit the exact mechanism requires further investigation.

Hemolymph sugar levels serve as cues for fine-tuning foraging behavior to maintain energy homeostasis (Simpson and Raubenheimer, 1993). Interestingly, RNAi of either Tret-1 or Tret-2 led to increased leaf consumption in this study. This suggests that the regulation of feeding behavior may be more complex than hemolymph trehalose levels alone, as Tret-1 knockdown decreased hemolymph trehalose levels, while Tret-2 knockdown increased it. It is likely that changes in trehalose levels within metabolically demanding tissues, such as muscle and the nervous system areas beyond the scope of this study, play a role in driving the observed increase in feeding behavior.

Since Tret-2 is primarily expressed in Malpighian tubules and plays a key role in transporting trehalose from hemolymph to these tubules, the uptake of trehalose into Malpighian tubules may contribute to the heat stress resilience associated with the aestivation state in CSFB. One of the primary functions of Malpighian tubules is osmoregulation, achieved through the absorption of water and solutes from the hemolymph (Farina et al., 2022). As the main hemolymph sugar in insects, trehalose may help establish the necessary energy and solute gradients to support this process. Osmoregulation is likely critical for heat stress resilience, as desiccation is a major cause of mortality under such conditions, with the excretory system contributing to overall water loss in insects (Koyama et al., 2021; Naseem et al., 2023).

### Conclusion

Two trehalose transporters were functionally characterized across the pre-aestivating, aestivating, and post-aestivating phases of the CSFB. In the fat body, trehalose peaks during the preparatory phase of aestivation and gradually decreases during the beetle’s ontogeny, whereas in the hemolymph and Malpighian tubules, trehalose levels peak mid-aestivation. This highlights the tightly regulated spatiotemporal dynamics of trehalose during aestivation. Tret-1 is responsible for transporting trehalose from the fat body into the hemolymph, while Tret-2 transports trehalose mainly into the Malpighian tubules. Additionally, a putative cross-talk fine-tunes the expression of each transporter to maintain trehalose homeostasis. Disrupting trehalose transport into the Malpighian tubules experimentally compromised heat stress resilience in aestivating beetles. These results provide a mechanistic understanding of spatiotemporal trehalose dynamics in the cabbage stem flea beetle and suggest trehalose transporters as potential targets for pest control via RNA interference.

## Acknowledgments

The authors thank Deutscher Akademischer Austauschdienst (DAAD) for research grants to Gözde Güney and Doga Cedden. The graphical abstract was created using BioRender.com, with assistance from Metin Cedden.

## Author Contributions

GG: Conceptualization, Writing - Original Draft, Investigation, Methodology, Formal analysis, Data Curation, Visualization. DC: Formal analysis, Methodology, Data Curation, Writing - Review & Editing. SS: Funding acquisition, Writing - Review & Editing. MR: Supervision, Funding acquisition, Writing - Review & Editing.

